# Nitric oxide synthase inhibition and biological sex define different cardiac responses to cardiometabolic stress in mature adult mice

**DOI:** 10.64898/2026.08.28.747952

**Authors:** June Sun, Alejandro Torres Riquileme, Joshua Hor, Daniel G. Donner, Helen Kiriazi, Shannon Walker, Simon T Bond, Natalie Mellet, Peter Meikle, Adam Parslow, Michael L.H. Huang, Monica Kanki, Brian G. Drew, Yow Keat Tham, Julie R. McMullen, Morag J. Young

**Author notes:** Correspondence: Professor Morag Young, PhD, Head, Cardiovascular Endocrinology laboratory group, Baker Heart and Diabetes Institute, 75 Commercial Road, Melbourne, Victoria 3004, Australia, PO Box 6492, Melbourne Victoria 3004, Australia. Equal contribution (June Sun, Alejandro Torres Riquileme).

## Abstract

**Background:** Heart failure (HF) with preserved ejection fraction (HFpEF) is a heterogeneous condition that accounts for more than half of all HF cases. It is a major cause of morbidity and mortality worldwide; however, preclinical models often do not capture the heterogeneity and sex-specific and cardiometabolic features observed in patients. We hypothesised that biological sex and the degree of nitric oxide synthase (NOS) inhibition would influence development of HFpEF-like versus HFrEF-like phenotypes.

**Methods and Results:** Male and female C57BL/6J mice (>24 weeks) were exposed to a high-fat diet (HFD) combined with low-dose (0.3 g/L; female only) or high-dose (0.5 g/L; male and female) NOS inhibition using N(ω)-nitro-L-arginine methyl ester (L-NAME) for 15 weeks. Female mice receiving low-dose L-NAME+HFD developed a HFpEF-like phenotype characterised by impaired diastolic function, exercise intolerance and preserved systolic function. Increasing NOS inhibition did not worsen diastolic dysfunction but induced inflammatory and stress-associated transcriptional pathways in female hearts. In contrast, male mice administered high-dose L-NAME+HFD developed hypertension, elevated ventricular pressures and impaired systolic function, consistent with a HFrEF-like phenotype. Despite different cardiac phenotypes, circulating lipidomic profiling revealed broadly conserved sphingolipid and phospholipid remodelling, with sex-specific regulation of phosphatidylinositol and lysophosphatidylcholine species. Transcriptomic analyses identified shared regulation of extracellular matrix, calcium-handling and metabolic pathways, whereas greater NOS inhibition was associated with transcriptional signatures related to inflammatory signalling, cellular stress responses and mitochondrial homeostasis.

**Conclusions:** Cardiometabolic stress does not produce a uniform HF phenotype. Instead, biological sex and the degree of NOS inhibition direct distinct functional and molecular remodelling trajectories, identifying HFpEF-like dysfunction as one of several potential cardiac responses to cardiometabolic injury in male and female mice.

## Introduction

Heart failure (HF) with preserved ejection fraction (HFpEF) now accounts for more than half of all HF cases and is a major cause of morbidity and mortality worldwide.^1 2^ Despite its increasing prevalence, effective therapies for HFpEF remain limited, in part because HFpEF represents a heterogeneous syndrome induced by multiple pathophysiological processes rather than a single disease entity.^3^ HFpEF is strongly associated with ageing, obesity, hypertension, metabolic dysfunction and systemic inflammation, all of which contribute to impaired ventricular relaxation, increased myocardial stiffness and exercise intolerance.^4^ HF with reduced ejection fraction (HFrEF), the biological pathways that initiate and sustain HFpEF remain incompletely understood.

Women account for more than half of HFpEF cases and exhibit a greater burden of obesity, metabolic dysfunction and vascular stiffening, particularly after menopause. ^5,6,7^ Ageing also plays a central role, with the incidence of disease increasing markedly in post-menopausal women,^8,9^ suggesting that the interaction between biological sex, ageing and cardiometabolic stress is a critical determinant of susceptibility to developing HFpEF.^10^ However, the molecular pathways linking metabolic dysfunction, endothelial impairment, and age-related loss of cardiovascular reserve to the development of HFpEF remain incompletely defined.

Progress towards understanding the pathogenesis of HFpEF has been limited by a lack of preclinical models that accurately reproduce the complexity, heterogeneity and greater female prevalence of HFpEF among patients.^11,12^ Existing models often rely on pressure-overload, neurohormonal activation or acute manipulation of cardiac stiffness and often progress towards systolic dysfunction.^11,13^ Combined high-fat diet (HFD) feeding and nitric oxide synthase (NOS) inhibition using N(ω)-nitro-L-arginine methyl ester (L-NAME) has emerged as a widely used “two-hit” model that incorporates metabolic stress and NOS inhibition.^14^ While this model reproduces several features of HFpEF, most studies have been conducted in 8-week-old mice and cardiac effects have been reported predominantly in male mice, limiting insights into age- and sex-specific disease mechanisms.^11,15^ Many studies have also focused on establishing a robust HFpEF phenotype rather than examining the factors that result in heterogeneous cardiac responses to cardiometabolic injury.^16,17,18^

One unresolved question in HFpEF models is whether disease severity simply reflects the magnitude of cardiometabolic injury or whether distinct biological responses determine different forms of HF. Clinical studies indicate that patients with similar risk factors, including obesity, hypertension and metabolic dysfunction, may develop markedly different cardiovascular phenotypes ranging from compensated diastolic dysfunction to overt systolic failure.^19^ These observations suggest that HFpEF may not represent a universal intermediate stage of disease, but rather a distinct response to cardiometabolic stress. Identifying the factors that determine these different functional outcomes remains a critical barrier to understanding HFpEF pathogenesis and developing targeted therapies.

Reduced nitric oxide bioavailability is a central feature of HFpEF and contributes to endothelial dysfunction, vascular stiffness, inflammation and impaired myocardial relaxation.^4^ However, endothelial dysfunction may not simply amplify a common disease process.^12,20^ Previous studies from our group suggest that female cardiovascular tissues are more sensitive to nitric oxide inhibition through mechanisms involving sex hormones, vascular reserve and metabolic adaptation.^21^ These observations raise the possibility that similar levels of cardiometabolic stress may induce fundamentally different remodelling responses in male vs female hearts. Furthermore, different levels of NOS inhibition may activate distinct biological responses to cardiometabolic stress.^22^

We therefore hypothesised that cardiac responses to cardiometabolic stress and the degree of NOS inhibition would differ in male and female mice of over 6 months in age. To test this hypothesis, we investigated varying combinations of ageing, obesity and levels of NOS inhibition in male and female mice and integrated physiological, haemodynamic, lipidomic and transcriptomic analyses to identify mechanisms that separate tissue remodelling, HFpEF-like dysfunction and progressive HF.

## Methods

### Animal model

Sixty-five C57BL/6J mice were studied: 40 females aged approximately 27 weeks and 25 males aged approximately 24 weeks (Animal Resources Centre, Murdoch, WA, Australia). Cages of female mice were randomly assigned (RAND() function in Excel) to receive control diet (11% energy from lipids, n=10), HFD (60% energy from lipids) plus 0.3 g/L L-NAME (n=15) or HFD plus 0.5 g/L L-NAME (n=15). Male mice were assigned to control diet (n=10) or HFD plus 0.5 g/L L-NAME (n=15). The lower L-NAME dose was evaluated only in females because our previous data suggested greater female sensitivity to NOS inhibition.^23^ All animal procedures were approved by the AMREP Animal Ethics Committee B (E/8404/2022/B). Mice were housed under 12-hr light/dark cycles with access to food and water ad libitum. Mice were weighed on a weekly basis and monitored daily for general body condition and activity. Tail cuff blood pressure was measured at 5 and 15 weeks in prewarmed mice (37C) over 3 consecutive days (CODA® blood pressure monitor).

### Cardiac Function

Echocardiography was performed using a VisualSonics Vevo F2 and UHF57x transducer under anaesthesia [i.p. injection of ketamine (80 mg/kg), xylazine (8 mg/kg), and atropine (0.96 mg/kg)]. Fur was removed from the chest with depilatory cream, and mice were placed in the supine position on a heat pad (∼37°C). Left ventricular (LV) function and structure were evaluated using parasternal long-axis and Doppler echocardiography to determine LV ejection fraction (EF) and mass index, global longitudinal strain (GLS), peak early (E) and late (A) filling velocities. Two-dimensional transthoracic echocardiography was used to record heart rate, early (E) and late (A) peak diastolic filling velocities, LV EF, LV mass index, and annular blood flow velocities: E’-wave, A’-wave, and S’-wave and E wave deceleration rate (EWDR). Mice were given atipamezole hydrochloride (0.2 mg/kg) to aid recovery.^24^

### Exercise Tolerance

Exercise tolerance was determined at 5 and 15 weeks using a treadmill-based, incremental exercise test as previously described.^25^ Mice underwent a familiarisation period with an Exer-3/6 Treadmill (Columbus Instruments, Ohio, USA) over 3 days, which involved progressively increasing intensities and time spent on the treadmill. On the day of formal testing, mice were fasted for 2 hr before testing. The treadmill speed was set at 10 m/min, which was increased by 2 m/min every 3 minutes; time, speed and distance ran for each mouse at exhaustion (defined as unable to keep pace with the treadmill, and refusal to run with gentle prodding) were analysed.^25^

### Intraperitoneal Glucose Tolerance Test

Glucose tolerance was determined at 5 and 15 weeks as previously described.^26,27^ Prior to glucose tolerance test (ipGTT) mice were fasted for 4 hr and tail blood collected for lipidomic analysis prior to i.p. glucose injection. Blood glucose was assessed at baseline and then at 15-, 30-, 45-, 60-, 90-, and 120-minutes post injection (Accu-Chek Performa, Roche Diagnostics, Basel, Switzerland).

### Cardiac pressure catheterisation

LV contractility and arterial blood pressure were evaluated at the endpoint in a subset of mice under 1.7% isoflurane anaesthesia via invasive pressure catheterisation (SPR-671, Millar, Netherlands) coupled to a data acquisition system (LabChart, ADInstruments, New Zealand). The catheter was positioned in the ascending aorta to record systolic, diastolic, pulse, and mean arterial pressures, and then advanced into the left ventricle to determine maximum and minimum dP/dt, end-diastolic pressure, and the time constant of relaxation (Tau).^28^

### Plasma lipid extraction and analysis

Lipids were extracted from plasma in 1:1 butanol and methanol (BUME) and combined with an internal standard mix (ISTD) diluted 1:10 with the BUME to create the final extraction solvent. Samples were vortexed for 10 sec, sonicated for 60 minutes at 18-22°C and centrifuged at 13,000 rpm, 10 min. 90 μl of the supernatant was transferred for analysis of lipid content by liquid chromatography-electrospray ionisation-tandem mass spectrometry using an Agilent 1290 HPLC coupled to an Agilent 6495C triple quadrupole mass spectrometer as previously described.^29^ Chromatographic data were analysed using Mass Hunter Quant B10.0 where relative lipid abundances were calculated by relating each area under the chromatogram for each lipid species to the corresponding internal standard. Correction factors were applied to adjust for different response factors and assessed by PCA analysis. Phosphatidylcholine (PC) species were excluded from subsequent analyses because differences were predominantly driven by dietary lipid composition (plant-based chow versus lard-based high-fat diet) rather than disease-related remodelling. ^30^

### RNA Isolation and quantitative real-time PCR

Total RNA was extracted from LV tissue using TRIzol (Thermo Fisher Scientific, MA) and Turbo DNAse I (Thermo Fisher Scientific), with 330 ng of RNA reverse transcribed using LunaScript® RT SuperMix Kit (New England Biolabs, MA) according to manufacturer’s instructions. Quantitative real-time PCR (qPCR) was then performed using 500 ng cDNA and Promega GoTaq 2X qPCR master mix (Promega, WI) and QuantStudio 7 Flex Real-Time PCR system (Thermo Fisher Scientific). Gene-specific primers are listed in **Supplementary Table S1**. Expression was analysed using ΔΔCt method by normalised to Hprt (housekeeping gene) and control group. Data are presented as fold change (2^-ΔΔCt^) relative to control.

### RNA-sequencing

RNA was extracted from mouse whole heart tissues with TRIzol (Invitrogen) and assessed for RNA integrity (Agilent Bioanalyzer 2100 and RNA Nano Assay, Agilent Technologies, CA, USA). Samples with RNA integrity number ≥7.6 were submitted for library construction (Novel RNA-Seq pipeline Version 01/09/2021) and sequenced (NextSeq2000) to generate ∼29bp single-end reads (n=5-12 per group) (Medical Genomics Facility, Monash University, Victoria, Australia). ∼19 million single-end reads were generated, and Fastq files were processed using the NfCore/RNAseq (3.10.1) pipeline using the “--with_umi” function. Reads were aligned to the Mus musculus GRCm38 reference using STAR aligner and quantified using featureCounts producing the raw genes count matrix and various quality control metrics. Raw counts were analysed with the DEGUST web tool to visualise differential gene expression and classical multidimensional scaling (MDS) and mean-difference plots (MA plots). First step visualisation of differentially expressed genes (DEG) in DEGUST included genes with a p(false-discovery rate [FDR]) ≤ 0.05 and change in expression ≥ 2-fold (|log2FC|≥1) using the limma/voom model.^31^

DEG were also analysed using QIAGEN Ingenuity Pathway Analysis (IPA, QIAGEN Digital Insights, Hilden, Germany) to determine enriched canonical pathways, upstream regulators and biological functions. Core Analysis was performed using the Ingenuity Knowledge Base (Genes Only) as the reference dataset, including both direct and indirect experimentally observed interactions derived from human, mouse and rat studies. DEG (defined as p(FDR) ≤0.05 and |log2FC| ≥1) and their log2FC value were used for IPA Core Analysis. Statistical significance of pathways identified was determined using Fisher’s exact test. IPA activation z-scores were used to predict activation (z-score≥2) or inhibition (z-score≤−2) of canonical pathways, upstream regulators and downstream biological functions. Regulator Effects and Causal Network analyses identified potential mechanistic links between upstream signalling events and downstream biological responses. Functional categories and disease associations were ranked according to enrichment significance and predicted activity state.

### Collagen staining

4 μM paraffin sections of LV tissue were dewaxed and rehydrated stepwise by 5-minute incubations in graded ethanol followed by a 5 min tap water before incubation in 0.1% w/v picrosirius red (PSR) stain for 30 min. Samples were rinsed 3x in water and rapidly dehydrated stepwise through graded ethanol followed by 2 x 5 min xylene and mounted in dibutyl phthalate polystyrene xylene (DPX;Merck-Millipore, city, country). Histological image analysis was performed on the QuPath software (v0.5.1) using bright-field scans of section (40x magnification, Monash Histology Platform, Clayton, Australia). Tissue features to be analysed were defined by predetermined settings in QuPath (**Supplementary Tables S2A-E)**.^24^ The minimum object size and minimum hole size were entered to accurately outline the tissue while excluding any empty spaces. Once the tissue was outlined successfully, the tissue area within the annotation was calculated and colour deconvolution performed on the outlined tissue. The PSR-positive area within the annotation was calculated and data exported from QuPath.^32^

### Power calculation, sample size determination and exclusion criteria

The study was designed to detect biologically meaningful differences in cardiac function between treatment groups. Sample size estimates were informed by published studies using comparable HFD+L-NAME HFpEF models,^14,21,33^ in which major echocardiographic indices of diastolic dysfunction (including E-wave velocity, E′ velocity and E/E′ ratio) differed by approximately 20-100% between control and HFpEF groups and experimental cohorts typically comprised 10-15 animals per group. Based on these data and the anticipated variability associated with ageing, obesity and NOS inhibition, a minimum sample size of n=10 mice per group was estimated to provide 80% power (α=0.05) to detect a biologically meaningful 15-20% difference in primary echocardiographic endpoints, including E-wave velocity, E-wave deceleration rate and LV volume measurements. To accommodate expected attrition during functional, histological, lipidomic and transcriptomic analyses, HFD+L-NAME treatment groups were expanded to n=15 animals per group. Investigators performing echocardiography, pressure-volume analyses, histological quantification, lipidomic analyses and transcriptomic analyses were blinded to treatment allocation until completion of primary data analysis. Exclusion criteria were predefined and included technical failure of echocardiographic or pressure-volume measurements, inadequate tissue quality for histological analyses, RNA integrity number below the threshold required for RNA sequencing, and failure of lipidomic quality-control standards. No animals or samples were excluded based on treatment response or statistical outcome.

### Statistical Analyses

Statistical analyses were performed using GraphPad Prism 10 software. Data were assessed for normality using the Shapiro-Wilk test. Datasets that satisfied assumptions of normality were assessed for normality using the Shapiro-Wilk test. Normally distributed data were analysed using unpaired t-test or one-way ANOVA with Tukey’s Post hoc test, whereas non-normally distributed datasets were analysed by Mann-Whitney U test or Kruskal-Wallis test with Dunn’s post hoc test). Statistical significance was defined as *P* < 0.05. Data are presented as mean ± SEM. Statistical analyses for lipidomic datasets were conducted using R Studio,^34^ with unpaired multiple comparison t-tests with Benjamini-Hochberg corrections were applied, with data presented as mean ± SEM.

## Results

### HFD induces glucose intolerance and exercise intolerance in ageing male and female mice

Male (∼24 weeks old) and female (∼27 weeks old) C57BL/6J mice were administered a HFD plus high-dose L-NAME (both male and female) or low-dose L-NAME (female only given females are more sensitive to NOS inhibition) for 15 weeks. Four mice died during the study such that the final sample size was male control (n=10), female control (n=8), male 0.5 g/L L-NAME (n=15), female 0.5 g/L L-NAME (n=15), and female 0.3g/L L-NAME (n=13). All mice gained weight over the 15-week study (**Figure 1A**). Female mice receiving either low-dose or high-dose L-NAME+HFD developed impaired glucose tolerance at 15 weeks, whereas increased body weight was most evident in the low-dose group. (**Figure 1B)**. Reduced exercise tolerance (running distance) at 15 weeks relative to control mice was evident in all HFD+L-NAME treated male and female groups (**Figure 1B-C**, **Table 1**). Wet tissue weights for left and right ventricles were higher in females given 0.3 g/L L-NAME+HFD versus control, but not when normalised to body weight. Kidney weight normalised to body weight was lower in 0.3 g/L L-NAME+HFD treated females, while female and male mice given 0.5 g/L L-NAME+HFD showed no change in any tissue weights (**Table 1**).

**Figure 1.**
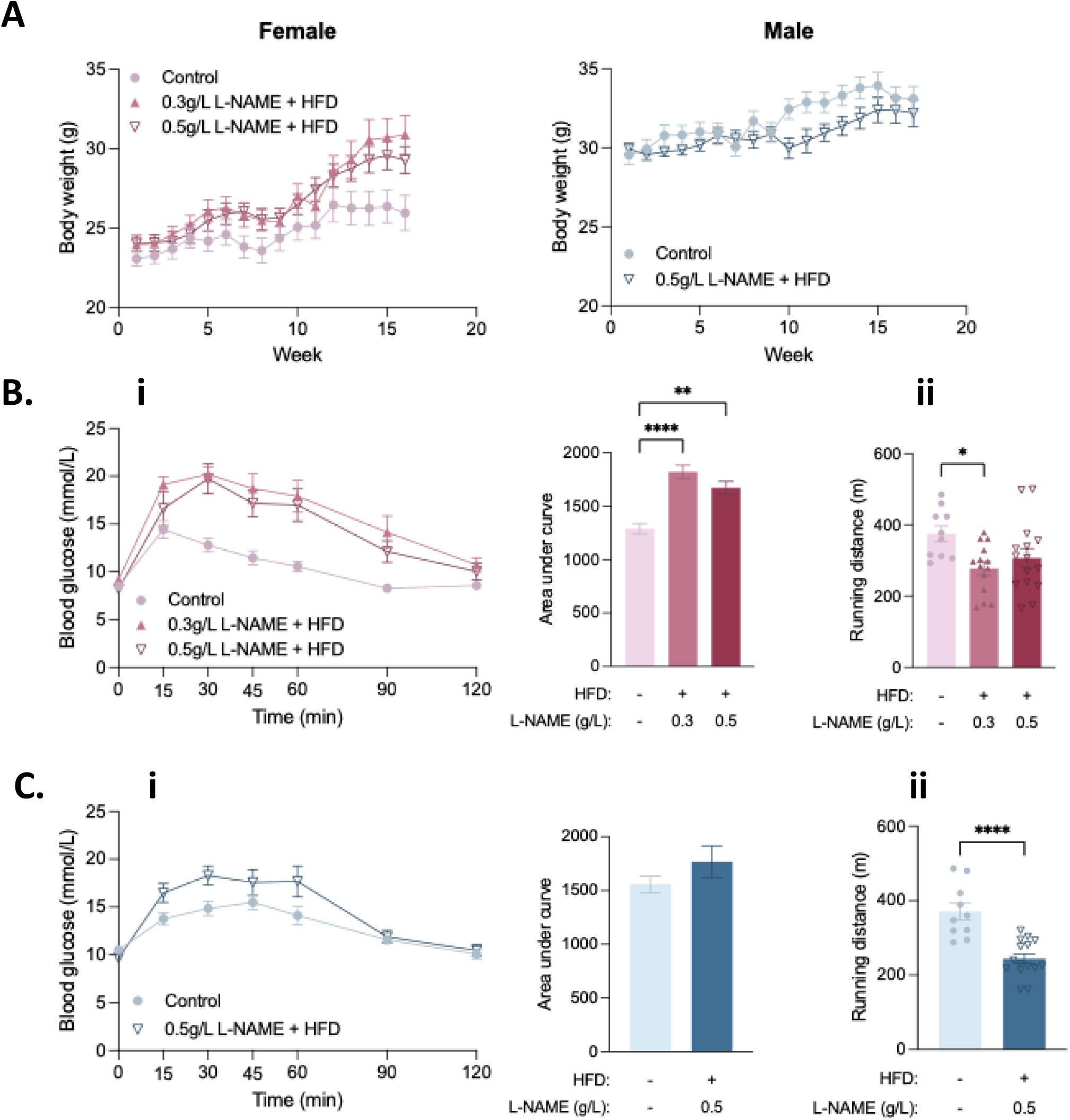
Cardiometabolic stress induces glucose intolerance and exercise intolerance. (A) Experimental study design showing treatment of male and female C57BL/6J mice with low- or high-dose L-NAME + HFD for 15 weeks. (B) i. Intraperitoneal glucose tolerance and ii. exercise tolerance in female mice at 15 weeks of treatment. (C) i. Intraperitoneal glucose tolerance and ii. exercise tolerance in male mice at 15 weeks. Data are presented as mean ± SEM and analysed by one way ANOVA (females) or unpaired T test (males). *P < 0.05, **P < 0.01 versus control.

**Table 1.** Exercise tolerance, glucose tolerance, body weight and tissue weights in female and male mice at 5 or 15 weeks of treatment: control diet, low-dose L-NAME (0.3 g/L) plus high-fat diet (HFD), or high-dose L-NAME (0.5 g/L) plus HFD. Parameters include treadmill exercise performance, intraperitoneal glucose tolerance test area under the curve (AUC), body weight and organ weights at 15 weeks post treatment. Female mice developed obesity, glucose intolerance and reduced exercise capacity following HFD+L-NAME treatment, whereas male mice demonstrated reduced exercise tolerance with less pronounced metabolic dysfunction. AUC: area under curve; BW: body weight; N/A: data not available. Data shown as mean ± SEM. \**P* < 0.05, \*\**P* < 0.01 versus control; ^*P* < 0.05 versus 0.3 g/L L-NAME + HFD.

|  | Control | 0.3 g/L L-NAME +<br>HFD | 0.5 g/L L-NAME +<br>HFD |
| --- | --- | --- | --- |
| Female | N=8-10 | N=13-15 | N=10-15 |
| *Average age (weeks)* |  |  |  |
| At start | 27.0±0.3 | 26.7±0.3 | 26.7±0.3 |
| *5-week treatment* |  |  |  |
| Running distance (m) | 369.2±32.1 | 365.7±16.7 | 325.5±45.2 |
| Running distance/BW<br>(m/g) | 15.4±1.5 | 14.3±0.9 | 13.1±1.9 |
| Blood glucose AUC | 1249±120 | 1429±96 | 1585±111 |
| Blood glucose/BW AUC | 52.3±5.5 | 55.3±4.1 | 62.9±4.7 |
| *15-week treatment* |  |  |  |
| Running distance (m) | 375.4±21.9 | 277.8±18.8* | 307.5±25.7 |
| Running distance/BW<br>(m/g) | 14.5±1.1 | 9.2±0.8** | 10.8±1.1* |
| Blood glucose AUC | 1289±48 | 1822±63** | 1672±60** |
| Blood glucose/BW AUC | 48.1±2.7 | 63.0±5.6 | 59.0±6.0 |
| *Wet tissue weights (mg)* |  |  |  |
| Left ventricle | 76.9±2.1 | 90.7±3.0** | 85.8±1.9 |
| Right ventricle | 20.7±2.6 | 22.9±0.6* | 19.8±1.0^ |
| Kidney | 147.5±6.9 | 143.3±4.7 | 147.8±4.7 |
| Lung | 187.1±10.1 | 213.0±11.2 | 186.0±11.4 |
| *BW (g)* |  |  |  |
| At start | 23.1±0.5 | 24.0±0.4 | 24.0±0.5 |
| At endpoint | 25.9±1.1 | 30.7±1.1* | 27.9±1.1 |
| *Tissue weight over endpoint*<br>*BW (mg/g)* |  |  |  |
| Left ventricle | 2.91±0.12 | 3.01±0.10 | 2.97±0.12 |
| Right ventricle | 0.78±0.09 | 0.76±0.03 | 0.68±0.04 |
| Kidney | 5.72±0.34 | 4.79±0.17* | 5.09±0.19 |
| Lung | 7.28±0.53 | 7.14±0.51 | 6.41±0.42 |
| Male | N=10 |  | N=15 |
| *Average age (weeks)* |  |  |  |
| At start | 24.0 | N/A | 24.0 |
| At endpoint | 40.0 | N/A | 40.0 |
| *5-week treatment* |  |  |  |
| Running distance (m) | 302.3±21.6 | N/A | 297.1±19.0** |
| Running distance/BW<br>(m/g) | 9.8±0.7 | N/A | 9.9±0.6 |
| Blood glucose AUC | 1294±91 | N/A | 1380±117 |
| Blood glucose/BW AUC | 42.1±3.5 | N/A | 45.6±3.6 |
| *15-week treatment* |  |  |  |
| Running distance (m) | 370.4±22.7 | N/A | 244.0±12.6 |
| Running distance/BW<br>(m/g) | 11.0±0.8 | N/A | 7.6±0.5** |
| Blood glucose AUC | 1556±77 | N/A | 1768±148 |
| Blood glucose/BW AUC | 46.0±2.3 | N/A | 54.5±4.2 |
| ***Raw tissue weights (mg)*** |  |  |  |
| Left ventricle | 114.0±3.2 | N/A | 112.8±3.1 |
| Right ventricle | 26.5±1.1 | N/A | 25.0±1.0 |
| Kidney | 188.0±3.7 | N/A | 190.2±4.3 |
| Lung | 180.8±13.7 | N/A | 206.5±11.1 |
| ***BW (g)*** |  |  |  |
| At start | 29.5±0.6 | N/A | 29.9±0.3 |
| At endpoint | 34.1±0.9 | N/A | 32.6±1.1 |
| ***Tissue weight over endpoint BW (mg/g)*** |  |  |  |
| Left ventricle | 3.45±0.09 | N/A | 3.54±0.14 |
| Right ventricle | 0.81±0.08 | N/A | 0.79±0.04 |
| Kidney | 5.70±0.14 | N/A | 5.97±0.22 |
| Lung | 5.53±0.50 | N/A | 6.56±0.46 |

### NOS inhibition preferentially induces diastolic dysfunction in female mice

Female mice given 0.3 g/L L-NAME+HFD showed evidence of early diastolic dysfunction with reduced peak E-wave velocity, E/A ratio, EWDR, IVRT, IVRTc, end-diastolic LV mass normalised to body weight (EDLVM/BW) and end-diastolic volume normalised to body weight (EDV/BW) compared with control animals **(Figure 2Ai-vi, Table 2A).** In contrast, females receiving 0.5 g/L L-NAME+HFD showed reduced E-wave velocity and EWDR only. Indicators of advanced diastolic dysfunction, including E/E′ ratio, E′ velocity, A′ velocity and deceleration time (dT), remained unchanged in both female treatment groups (**Table 2A**). Male mice given 0.5 g/L L-NAME+HFD showed a reduced peak E-wave velocity, peak A-wave velocity, EWDR and E′ velocity, but no change in E/A ratio, E/E′ ratio or ventricular mass indices versus control (**Figure 2Bi-vi, Table 2B**).

**Figure 2.**
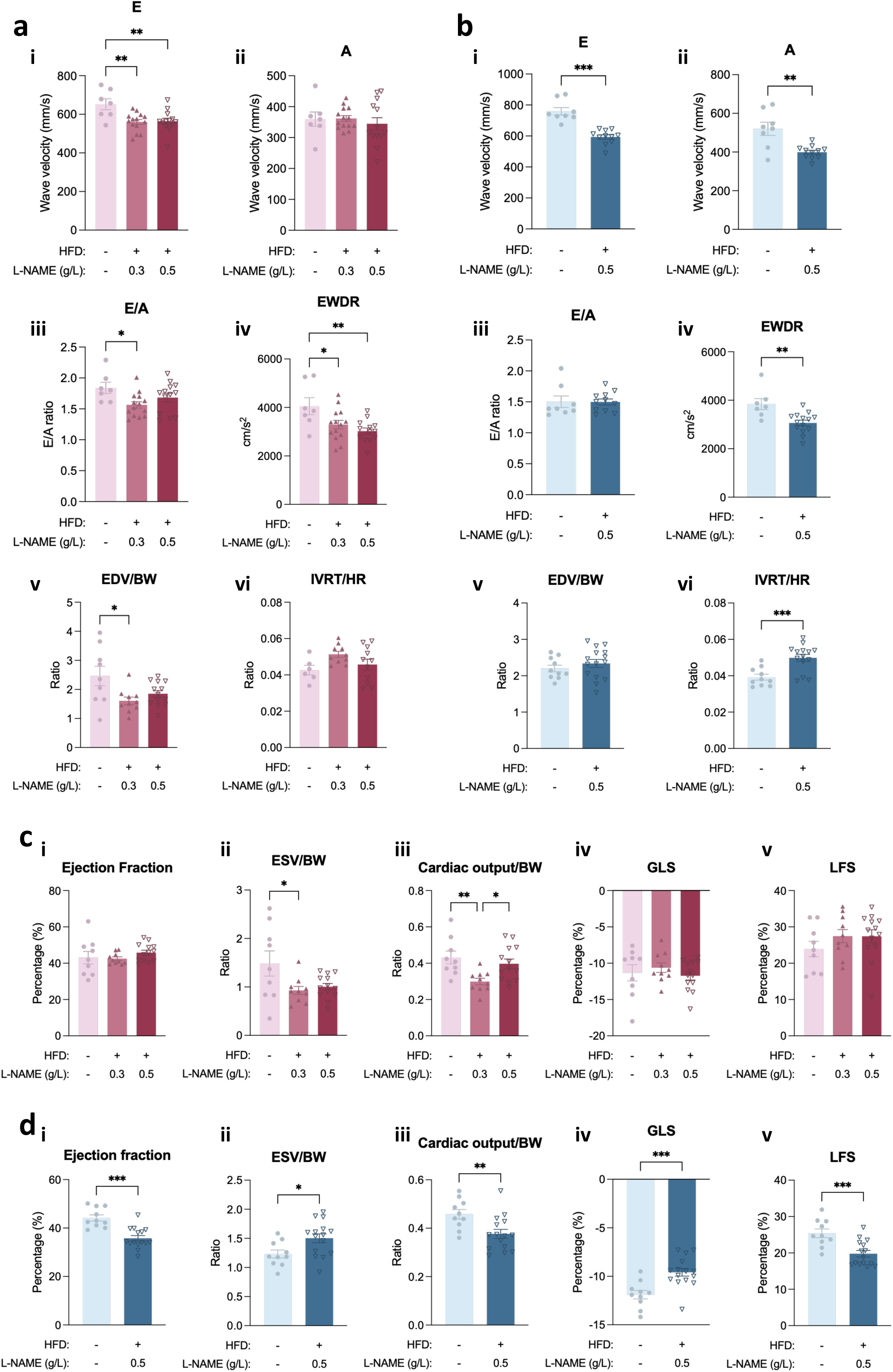
NOS inhibition plus cardiometabolic stress differentially affects cardiac function in female and male mice. (A) Echocardiographic indices of diastolic function in female mice following 15 weeks of treatment. (B) Echocardiographic indices of diastolic function in male mice following 15 weeks of treatment. (C) Echocardiographic indices of systolic function in female mice. (D) Echocardiographic indices of systolic function in male mice. Low-dose L-NAME induced a HFpEF-like phenotype in females characterised by impaired diastolic function with preserved systolic performance, whereas male mice exposed to HFD plus high-dose L-NAME developed systolic dysfunction. Data are presented as mean ± SEM and analysed by one way ANOVA or Kruskal-Wallis test for datasets not normally distributed, and unpaired T test as appropriate. \**P* < 0.05, \*\**P* < 0.01, \*\*\**P* < 0.001 versus control.

**Table 2A.** Echocardiographic assessment of cardiac structure and function in female mice. Echocardiographic measurements obtained after 15 weeks of treatment in female mice receiving control diet, 0.3 g/L L-NAME plus high-fat diet (HFD), or 0.5 g/L L-NAME plus HFD. Parameters include heart rate, indices of diastolic function, ventricular dimensions, ventricular mass, systolic function and cardiac output. BW: body weight; DT: deceleration time; DTc: deceleration time corrected for heart rate; EDLVM: end-diastolic left ventricular mass; EDV: end-diastolic volume; ESV: end-systolic volume; EWDR: E-wave deceleration rate; HR: heart rate; IVRT: isovolumic relaxation time; IVRTc: isovolumic relaxation time corrected for heart rate. Data shown as mean ± SEM. *P < 0.05, **P < 0.01 compared to control; ^P < 0.05 compared to 0.3 g/L L-NAME + HFD.

|  | **Control** | **0.3 g/L L-NAME+HFD** | **0.5 g/L L-NAME + HFD** |
| --- | --- | --- | --- |
|  | N=7-8 | N=10-14 | N=13-14 |
| HR (bpm) | 442±18 | 426±6 | 420±11 |
| IVRT (ms) | 18.86±0.82 | 22.70±0.50** | 21.43±0.81 |
| IVRT<sub>c</sub> (ms) | 50.22±2.07 | 60.39±1.26** | 55.99±1.91 |
| E (mm/s) | 652.14±28.49 | 560.56±12.73** | 563.82±15.72** |
| A (mm/s) | 359.52±23.03 | 361.77±9.35 | 344.62±19.85 |
| E' (mm/s) | 23.62±2.05 | 21.85±1.81 | 20.60±1.56 |
| A' (mm/s) | 17.39±1.10 | 16.73±0.68 | 17.11±0.94 |
| E/A | 1.84±0.09 | 1.57±0.05* | 1.68±0.07 |
| E'/A' | 1.29±0.10 | 1.31±0.09 | 1.22±0.07 |
| E/E' | 28.47±2.06 | 28.88±3.24 | 29.32±2.26 |
| DT (ms) | 16.63±1.13 | 17.49±0.73 | 19.09±0.98 |
| DT<sub>c</sub> (ms) | 44.34±3.25 | 46.65±2.13 | 50.11±2.76 |
| EWDR (cm/s<sup>2</sup>) | 4053±349 | 3291±175* | 3023±127** |
| EDLVM (mg) | 64.82±5.91 | 56.12±1.86 | 61.05±2.15 |
| EDLVM/BW (mg/g) | 2.55±0.31 | 1.84±0.09* | 2.26±0.10^ |
| EDV (μl) | 62.90±7.72 | 48.93±2.96 | 49.60±1.89 |
| ESV (μl) | 37.56±6.14 | 28.19±1.91 | 26.85±1.21 |
| EDV/BW (μl/g) | 2.46±0.33 | 1.61±0.13* | 1.85±0.12 |
| ESV/BW (μl/g) | 1.48±0.26 | 0.93±0.08* | 1.00±0.07 |
| Stroke volume (μl) | 25.34±1.86 | 20.76±1.19 | 22.76±1.11 |
| Stroke volume/BW (μl/g) | 0.979±0.077 | 0.680±0.048** | 0.848±0.056 |
| Ejection fraction (%) | 43±3 | 43±1 | 46±1 |
| Global longitudinal strain (%) | -11.3±1.1 | -10.6±0.6 | -11.7±0.6 |
| Cardiac output (ml/min) | 11.17±0.78 | 9.18±0.50 | 10.77±0.69 |
| Cardiac output/BW | 0.432±0.035 | 0.299±0.018** | 0.397±0.027^ |

### High-dose L-NAME plus HFD induces systolic dysfunction in males but not females

Female mice maintained preserved systolic performance following cardiometabolic stress. EF and GLS remained unchanged in both female groups given L-NAME+HFD (**Figure 2Ci-v, Table 2A**). Body weight-adjusted cardiac output and end-systolic volume indices were lower in females receiving 0.3 g/L L-NAME+HFD relative to the control and 0.5 g/L L-NAME+HFD (not significant for ESV/BW) groups, but overall systolic function was preserved in female mice given the lower dose of L-NAME (**Figure 2Cii-iii, Table 2A**). In contrast, male mice exposed to 0.5 g/L L-NAME+HFD developed overt systolic dysfunction, which was characterised by reductions in EF, cardiac output/body weight and longitudinal fractional shortening (LFS), together with increased end-systolic volume/body weight. GLS values became less negative, consistent with impaired myocardial deformation and reduced longitudinal contraction (**Figure 2Di-v, Table 2B**).

**Table 2B.**
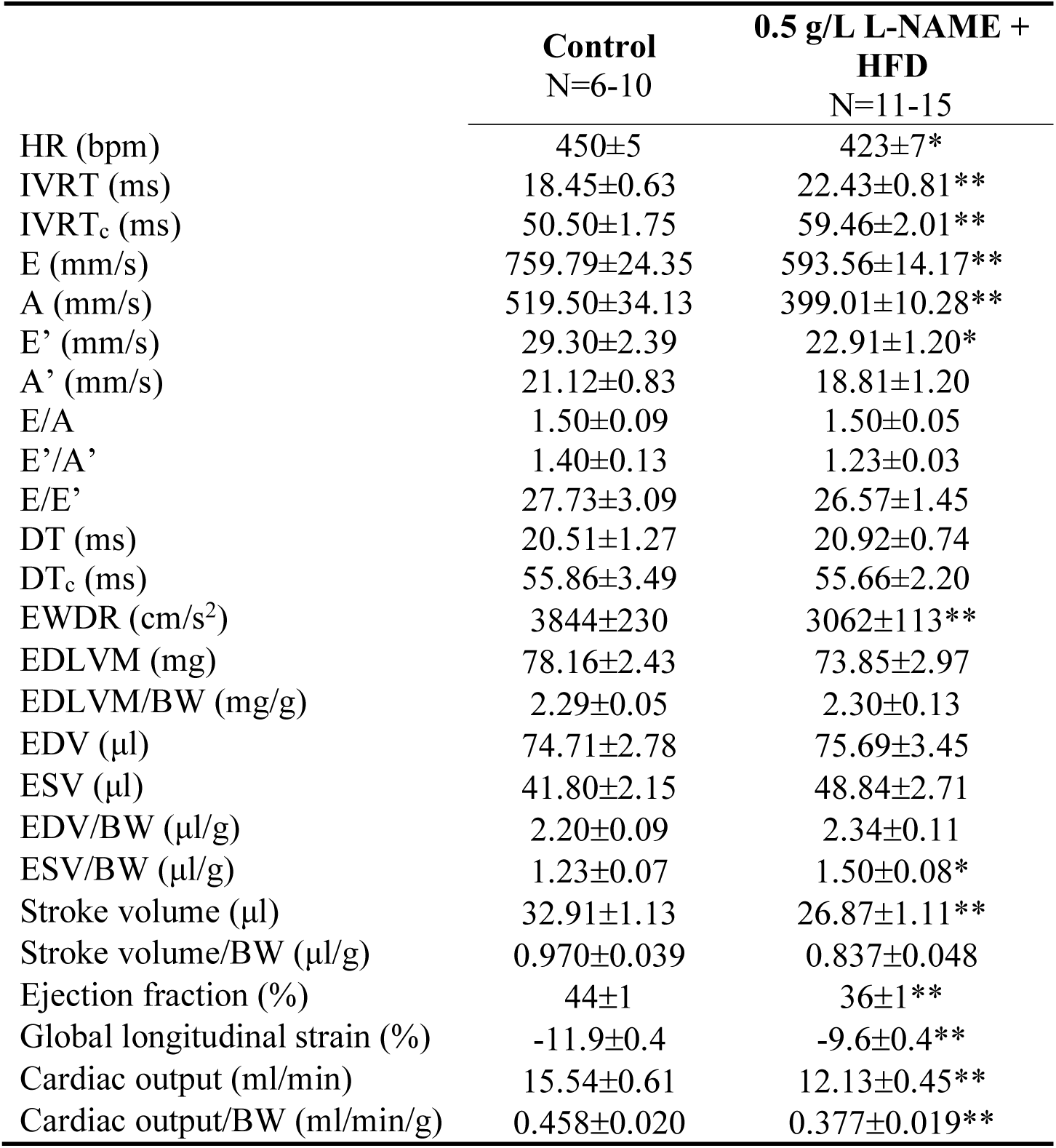
Echocardiographic assessment of cardiac structure and function in male mice. Echocardiographic measurements obtained after 15 weeks of treatment in male mice receiving control diet or 0.5 g/L L-NAME plus HFD. Parameters include heart rate, indices of diastolic function, ventricular dimensions, ventricular mass, systolic function and cardiac output. BW: body weight; DT: deceleration time; DTc: deceleration time corrected for heart rate; EDLVM: end-diastolic left ventricular mass; EDV: end-diastolic volume; ESV: end-systolic volume; EWDR: E-wave deceleration rate; HR: heart rate; IVRT: isovolumic relaxation time; IVRTc: isovolumic relaxation time corrected for heart rate. Data are presented as mean ± SEM. \**P* < 0.05, \*\**P* < 0.01 compared to control.

### Hypertension and haemodynamic dysfunction are selectively induced in male mice

To determine whether differential haemodynamic responses contributed to the observed cardiac phenotypes, blood pressure measurements for all mice and invasive pressure-volume analyses for a subset of hearts from control and 0.5 g/L L-NAME+HFD treated male and female mice were performed. The latter analyses compared cardiac haemodynamic profiles directly between male and female mice. Male mice receiving 0.5 g/L L-NAME+HFD developed sustained elevations in systolic, diastolic and mean blood pressure at both 5 and 15 weeks (**Figure 3B, D and F**). Consistent with these data, invasive haemodynamic assessment demonstrated increased systolic, diastolic and mean arterial pressures, together with elevated LV pressures and reduced contractility indices in male mice (**Figure 3E-F, Supplementary Table S3**). In contrast, in female mice tail-cuff blood pressure in mice receiving 0.3 or 0.5 g/L L-NAME+HFD was not different to control (**Figure 3A, C and E**). Mean arterial pressure was preserved (**Figure 3**), while ventricular relaxation indices remained unchanged (**Supplementary Table S3).**

**Figure 3.**
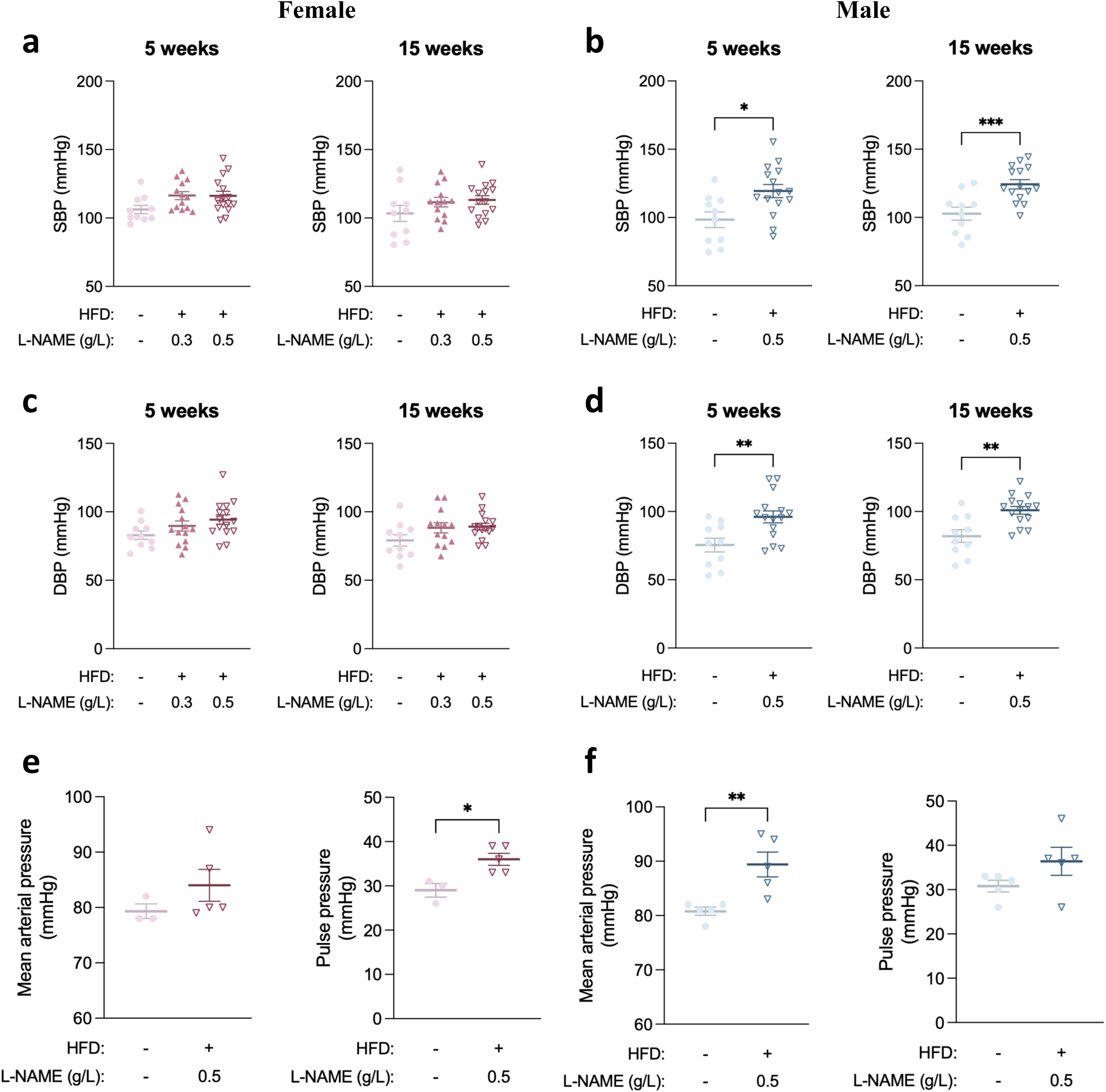
Haemodynamic responses to cardiometabolic stress differ between male and female mice. Tail-cuff blood pressure and invasive pressure catheter analyses at 5 or 15 weeks of treatment. (A-B) Blood pressure measurements obtained during the treatment period. (C-D) Left ventricular pressure catheter parameters. (E-F) Aortic pressure catheter parameters. Male mice developed hypertension and impaired ventricular function, whereas female mice exhibited relatively preserved LV haemodynamic regulation with selective changes in pulse pressure and heart rate. Data are presented as mean ± SEM and analysed by one way ANOVA or Mann-Whitney test for datasets not normally distributed, and unpaired T test as appropriate. *P < 0.05, **P < 0.01, ***P<0.001 versus control.

### Functional impairment occurs in the absence of overt myocardial fibrosis

We next quantified cardiac collagen deposition using picrosirius red staining and automated image analysis. Neither male nor female treatment groups exhibited a change in interstitial fibrosis despite substantial physiological and molecular remodelling. Although baseline collagen content was higher in female hearts compared with males, cardiometabolic stress did not increase collagen deposition in either sex at this time point (**Supplementary Figure S1**).

### Serum lipidomic profiling identifies shared and sex-specific metabolic responses to cardiometabolic stress

PCA analysis of plasma lipidome profiles demonstrated clear separation between treatment and control groups, particularly at the 15-week endpoint (**Figure 4A/4B)** for males and females, and at 5-week in the males (**Supplementary Figure S2)**, indicating substantial dysregulation of the circulating lipidome following L-NAME+HFD administration.

**Figure 4.**
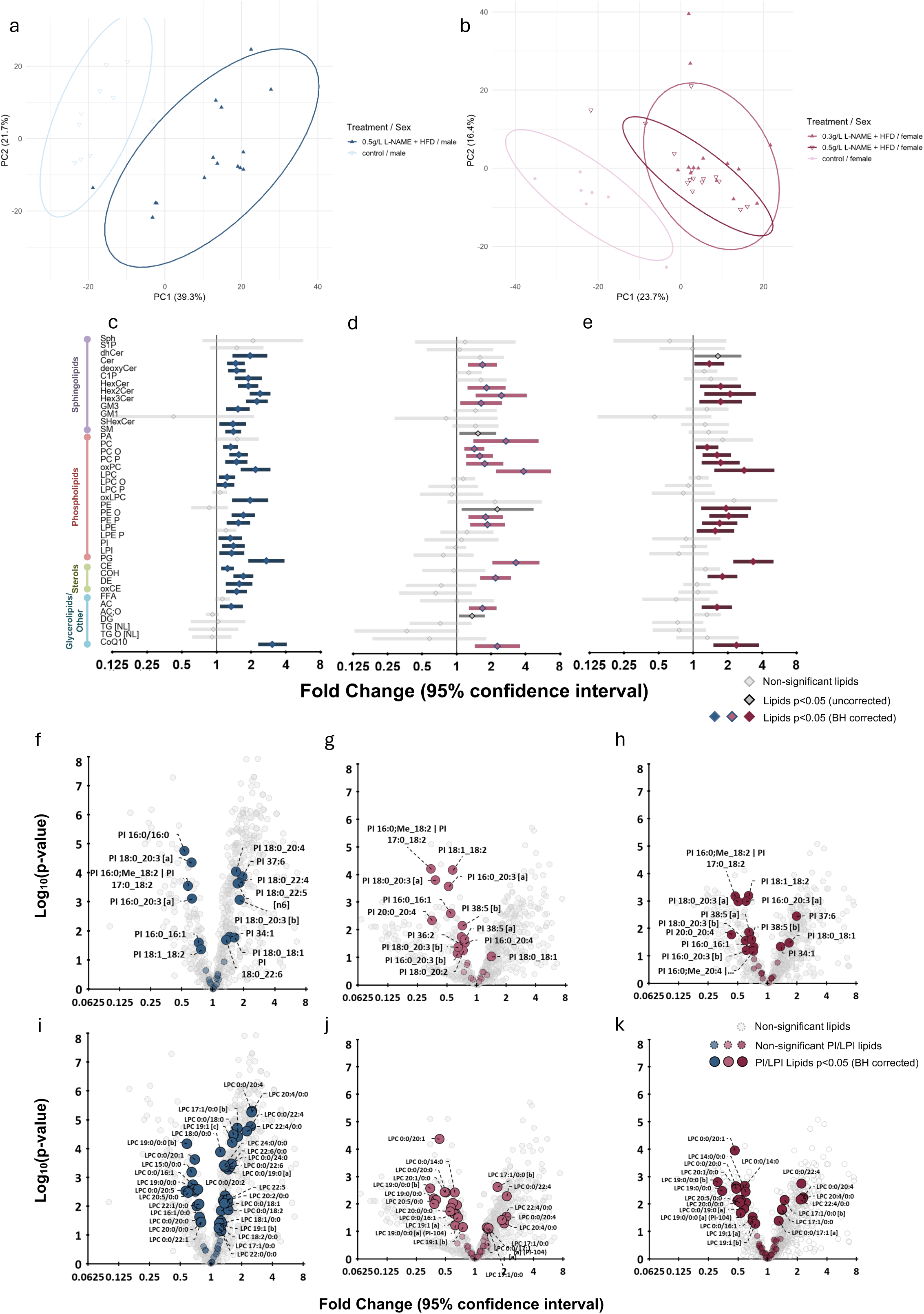
Cardiometabolic stress induces conserved lipidomic remodelling in male and female mice. (A-B) Principal component analysis of circulating lipid profiles. (C-E) Fold-change analysis of major lipid classes at study endpoint. (F-I) Regulation of sphingolipid, phospholipid, sterol and glycerol pathways. (J) Differentially regulated lipid species. Treatment induced broad remodelling of circulating sphingolipid and phospholipid classes in both sexes despite divergent cardiac phenotypes. Data are presented as fold-change relative to control with 95% confidence intervals.

Despite distinct clustering by sex and treatment, several lipid classes were regulated in a similar direction in both sexes, including the sphingolipids, phospholipids, ceramides, hexosylceramides, phosphatidylglycerols and plasmalogens, while diacylglyceride and triacylglyceride classes were mostly unchanged. Both sexes demonstrated an increase in multiple sphingolipid and phospholipid classes, including ceramides (Cer), monohexosylceramides (HexCer) and dihexosylceramides (Hex2Cer), phosphatidylglycerols, plasmalogens, and several lysophospholipid classes, while diacylglycerol and triacylglycerol classes remained largely unchanged and selected neutral lipid classes, including CoQ10 and acylcarnitines, were increased **(Figures 4C-E, Supplementary Figures S2)**.

Two key lipid classes, phosphoinositols (PIs) and lysophosphatidylcholines (LPCs), demonstrated sex-specific responses to HFD+L-NAME treatment (**Figure 4F-K**). In serum from male HFD+L-NAME mice, individual PI species showed variable regulation, with species containing longer-chain fatty acyl groups (20:4, 22:4, 22:5 and 22:6) generally increased (**Figure 4F**). In contrast, these increases were not observed in female HFD+L-NAME mice, in which most regulated PI species were decreased (**Figure 4G-H**). Similarly, LPC species demonstrated sex-specific regulation. Although male HFD+L-NAME mice exhibited both increases and decreases in individual LPC species, a greater number of LPC species were increased compared with female HFD+L-NAME groups, in which the relatively few regulated LPC species were almost exclusively those containing 17:1, 20:4 and 22:4 fatty acyl chains (**Figure 4I-K**).

### The metabolic stress responses differ between male and female hearts

To determine whether the distinct cardiac phenotypes were associated with sex-specific molecular responses, expression of genes linked to cellular stress, cardiac injury, inflammation and extracellular matrix remodelling was assessed by RT-qPCR. In male left ventricular tissue, high-dose L-NAME+HFD was associated with increased expression of the integrated stress response genes *Atf4, Atf6* and *Chop*, together with increased expression of the cardiac stress marker *Nppa* (**Figure 5**). Expression of Myh2 was also down regulated, whereas *Mmp9* and *Tlr7* mRNA levels were not different in males given L-NAME+HFD. In contrast, female hearts exhibited a more limited transcriptional response to treatment. Compared with controls, female mice given low dose, but not high dose, L-NAME+HFD showed reduced expression of *Mmp9* and increased expression of *Nppb*, and both treatment groups showed increased expression of *Tlr7*, all other genes tested (*Atf4, Atf6 or Chop*) were not different (**Figure 5**).

**Figure 5.**
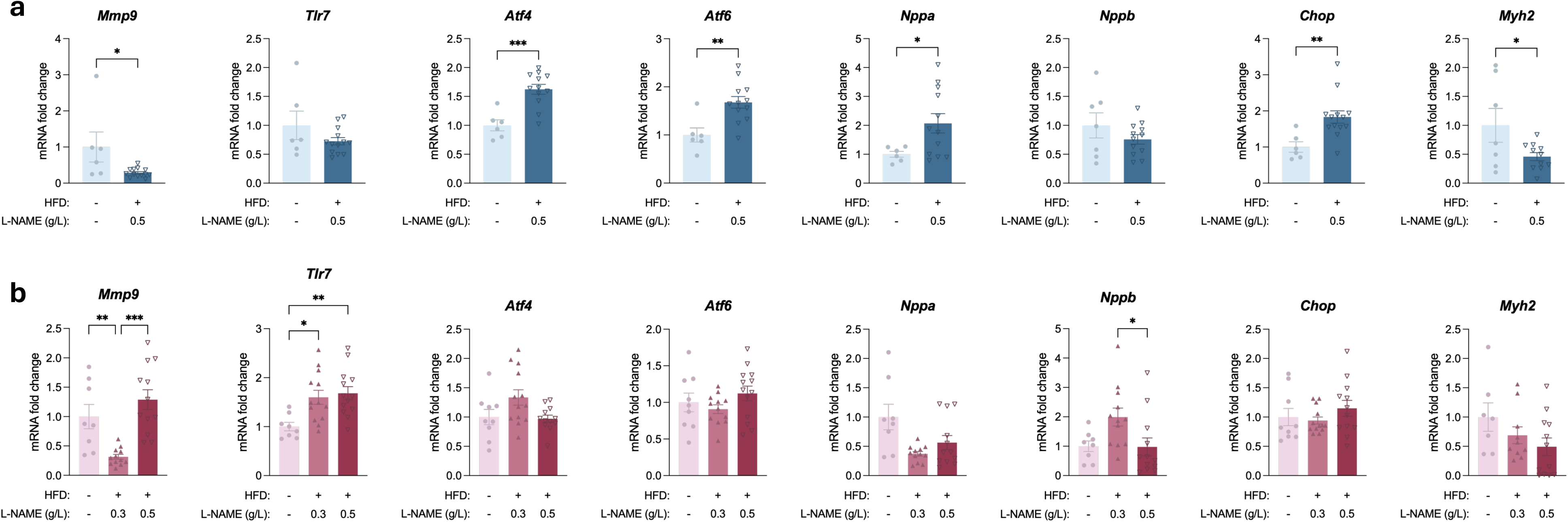
Molecular responses to cardiometabolic stress differ between male and female hearts. Quantitative PCR analysis of genes associated with cellular stress, inflammation, extracellular matrix remodelling and cardiac injury following 15 weeks of treatment. Male hearts demonstrated induction of stress- and injury-associated genes, whereas female hearts exhibited a more restricted transcriptional response. Data are presented as relative gene expression normalised to housekeeping genes and expressed relative to control, mean ± SEM and analysed by one way ANOVA (female, Mmp9 and Tlr7), or Kruskal-Wallis (female, Nppb) test for datasets not normally distributed, or unpaired T test (male, Atf6, Atf4, Nppa, Myh2) or Mann-Whitney (male, Mmp9 and Chop) as appropriate. *P < 0.05, **P < 0.01, *** P<0.001, **** P<0.0001 versus control.

### Distinct transcriptomic pathways underlie adaptive and maladaptive tissue remodelling in female hearts

Transcriptional responses differed substantially between the low- and high-dose L-NAME+HFD groups (**Figure 6A-C**). In low-dose L-NAME+HFD hearts, canonical pathway analysis identified enrichment of pathways related to extracellular matrix organisation, mitochondrial biology and dilated cardiomyopathy-associated signalling (**Figure 6D-H**). Extracellular matrix remodelling was characterised by increased expression of genes encoding collagens and matricellular proteins, including *Col3a1, Col4a1, Col4a2, Col6a1, Serpinh1, Sparc, Fstl1, Dpt, Lgals1* and *Serping1* (**Figure 6D-E**). Additional regulated pathways included mechanotransduction and myocardial contractile function, with reduced expression of *Dst, Dsp, Cmya5, Ttn, Tnni3* and *Nebl* (**Figure 6D-E**). Both treatment groups demonstrated coordinated regulation of calcium-handling genes including *Ryr2, Trdn, Hrc, Kcnip2, Kcnd2* and *Trpm4*, suggesting altered excitation-contraction coupling as a common response to cardiometabolic stress. The low-dose group also exhibited altered expression of genes involved in mitochondrial quality control and metabolism, including *Pink1, Mfn2, Coq10a, Ppm1k, Ivd, Oxct1, Atp5a1, Ogdh* and *mt-Rnr1* (**Figure 6F-H**).

**Figure 6.**
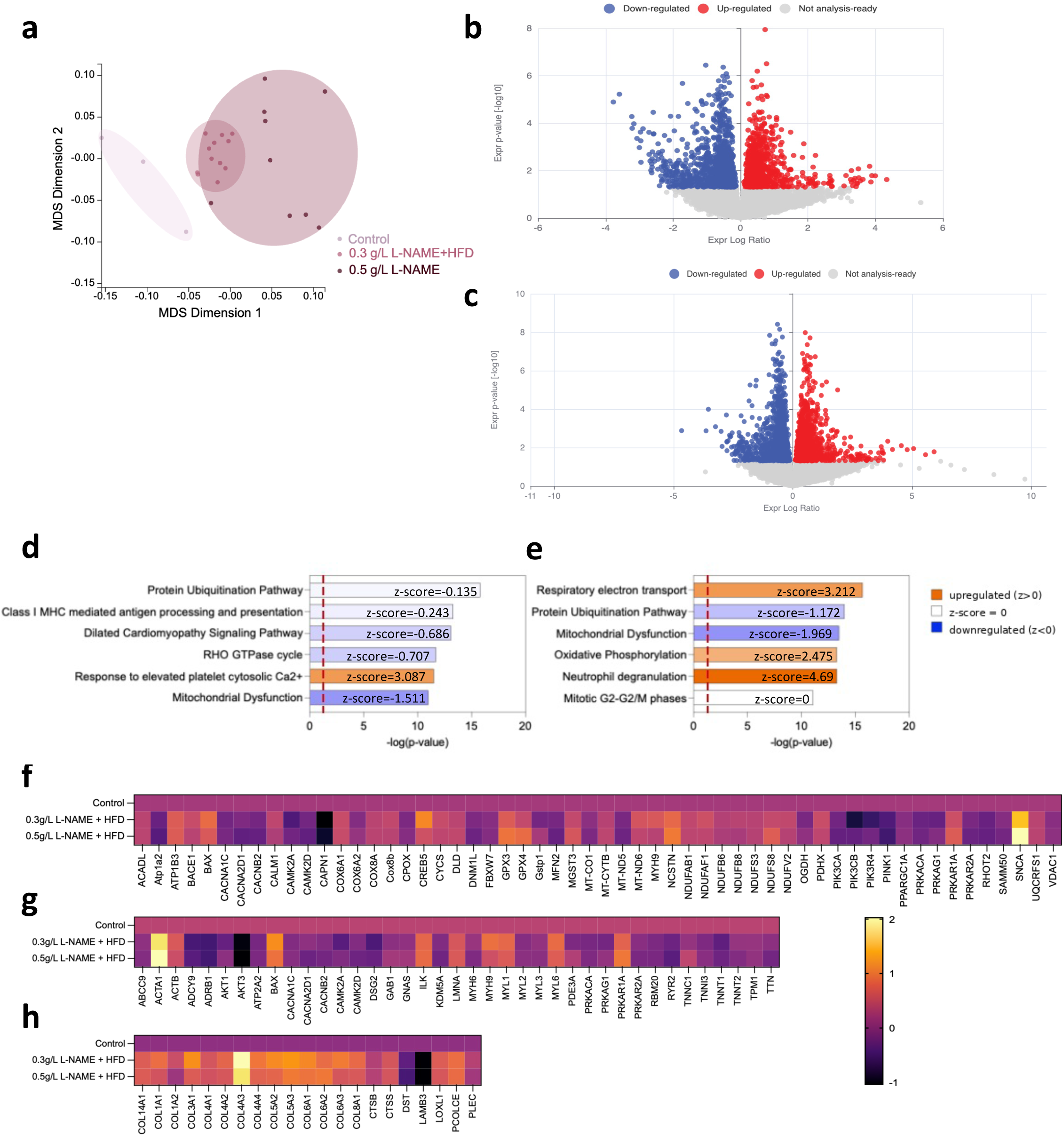
0.5 g/L L-NAME+HFD inhibition shifts cardiac remodelling from adaptive metabolic responses towards transcription of inflammatory and injury-associated pathways. RNA sequencing was performed on the LV from female mice at 15 weeks of treatment with control diet, low-dose L-NAME (0.3 g/L) plus high-fat diet (HFD), or high-dose L-NAME (0.5 g/L) plus HFD. (A) Multidimensional scaling analysis demonstrating distinct clustering of control, low-dose L-NAME/HFD and high-dose L-NAME/HFD groups, indicating that the higher dose of L-NAME in female mice produced distinct transcriptional responses. (B-C) Volcano plots showing differentially expressed genes in low-dose L-NAME/HFD versus control and high-dose L-NAME/HFD versus control comparisons. Differentially expressed genes were defined as FDR-adjusted P < 0.05 and |log2 fold-change| ≥ 1. (D) Enrichment of mitochondrial pathways demonstrating regulation of genes involved in mitochondrial maintenance, oxidative stress responses and cellular energetics. (E) Cardiomyocyte structure and function pathways, including genes associated with excitation-contraction coupling, calcium handling and cardiomyopathy-related signalling. (F) Extracellular matrix organisation and collagen assembly pathways demonstrating activation of structural remodelling programmes. (G-H) Additional enriched biological pathways illustrating divergence between adaptive remodelling and injury-associated responses with increasing endothelial stress.

High-dose L-NAME+HFD induced a distinct transcriptomic profile characterised by predicted activation of oxidative phosphorylation (z = 2.475) and respiratory electron transport pathways (z = 3.212), together with enrichment of inflammatory signalling pathways (z = 4.69) (**Figure 6F-H**). These changes occurred alongside altered expression of genes involved in mitochondrial maintenance and homeostasis, including *Pink1, Mfn2, Samm50, Oxct1, Coq10a, Atp5a1, Lrpprc, mt-Nd5, mt-Rnr1* and *mt-Rnr2*, as well as increased expression of inflammatory and stress-associated genes including *Ifitm3, Lyz2, Anxa1, Cd9, Cd81, Gpx3, Gpx4, Bnip3, Depp1, Cdkn1a* and *H2-D1* (**Figure 6F-H; Supplementary Tables S4-S5**). We also observed altered expression of circadian-clock genes (*Per3, Cry2, Dbp, Tef* and *Hlf*) and corticosteroid-responsive genes (*Sgk1, Fkbp5, Tsc22d3* and *Pdk4*). As circadian and corticosteroid signalling regulate myocardial metabolism, mitochondrial function and inflammatory responses, these pathways may contribute to cardiac adaptation during chronic cardiometabolic stress. (**Supplementary Tables S4-S5**).

## Discussion

The major finding of this study is that similar cardiometabolic stressors produced fundamentally different HF phenotypes in mature, but not senescent, adult male and female mice. Increasing L-NAME exposure did not intensify the HFpEF phenotype but instead was associated with activation of inflammatory, mitochondrial and injury-associated transcriptional pathways. By combining obesity, different levels of NOS inhibition and mice of older age (>6 months at study commencement), we showed that HF phenotype is not determined solely by the extent of cardiometabolic injury, but by sex-specific responses to endothelial and metabolic stress. Female mice exposed to HFD and low-dose L-NAME developed the most consistent HFpEF-like phenotype that was characterised by exercise intolerance, impaired glucose handling, abnormalities in diastolic function and preserved systolic performance. In contrast, male mice exposed to similar cardiometabolic stressors progressed towards hypertension and systolic dysfunction (i.e. HFrEF). These findings raise the possibility that HFpEF-like remodelling occurs within a limited range of NOS inhibition rather than increasing linearly with higher L-NAME exposure.

HFpEF is increasingly recognised as a heterogeneous syndrome rather than a single disease.^17,35,36^ Clinical classification approaches now recognise multiple HFpEF endotypes driven by differing contributions of obesity, metabolic dysfunction, endothelial dysfunction, inflammation, ageing and renal disease. Similarly, recent evaluations of preclinical HFpEF models have concluded that no single model reproduces the full complexity of the human syndrome and that female animals remain markedly under represented despite women comprising the largest proportion of HFpEF patients.^15,16^ The present study addresses several of these limitations by incorporating both sexes, using older animals than many previous studies, and directly examining how NOS inhibition modifies the myocardial response.^16^ Rather than reproducing a single HFpEF phenotype, our data show that biological sex and NOS inhibition influences the development myocardial pathology and distinct forms of HF.

One unexpected finding was that female mice exposed to low-dose L-NAME developed a more clinically relevant HFpEF-like phenotype than females receiving the higher L-NAME dose. Low-dose L-NAME was associated with impaired ventricular filling, reduced exercise capacity and metabolic dysfunction while preserving systolic performance. In contrast, high-dose L-NAME did not enhance diastolic dysfunction and instead promoted activation of inflammatory, oxidative stress and mitochondrial stress pathways. These data suggest that low-dose and high-dose L-NAME exposure activate distinct physiological and molecular responses to cardiometabolic stress where the lower dose group produced a phenotype more consistent with HFpEF-like remodelling, while the higher dose group resulted in activation of inflammatory and stress-related transcriptional pathways.^36,37^

A major strength of this study is the direct comparison of male and female responses and that we investigated mice that were of at least 6 months in age, i.e. mature but not senescent. Despite experiencing similar systemic metabolic stress, female mice maintained preserved systolic function while males developed hypertension, elevated ventricular loading conditions, reduced EF, impaired myocardial deformation and reduced cardiac output. These observations are relevant clinically because women account for the majority of HFpEF cases, yet most preclinical HFpEF studies continue to use young male animals.^16^ The present findings suggest that sex does not simply influence the severity of disease but fundamentally alters the biological response to cardiometabolic injury. Differences in endothelial adaptation to nitric oxide inhibition, vascular reserve, myocardial energetics, mitochondrial resilience and inflammatory signalling are likely to contribute to these variable outcomes.

Transcriptomic analyses provided important mechanistic insight into these different phenotypes and suggest that increasing NOS inhibition does not simply worsen existing cardiac abnormalities but changes the myocardial response to reduced energetic resilience and activation of injury-associated pathways.^38,39^ Increasing evidence suggests that impaired metabolic flexibility is a central feature of HFpEF pathogenesis.^40^ The healthy myocardium maintains substantial flexibility in substrate selection and can adapt fuel utilisation according to metabolic demand.^41,42^ In contrast, HFpEF is increasingly recognised as a disorder characterised by impaired metabolic adaptation, altered fatty acid utilisation, mitochondrial dysfunction and energetic insufficiency.^41,43^ Recent studies have demonstrated that disruption of NAD+-dependent signalling, mitochondrial quality-control pathways and ketone metabolism contributes directly to development of HFpEF, while restoration of these pathways improves cardiac function and reverses diastolic dysfunction.^44 45^ Consistent with these studies, we identified coordinated regulation of genes involved in mitochondrial maintenance, oxidative stress responses and energetic adaptation, particularly in female mice exposed to high-dose L-NAME. Recent studies increasingly implicate impaired metabolic flexibility and mitochondrial dysfunction as important contributors to HFpEF pathogenesis, suggesting that energetic reserve influences the tissue pathogenesis, independently of haemodynamic load or fibrosis.^46^ The present study thus identified regulation of pathways required for mitochondrial maintenance, oxidative stress responses, calcium handling and lipid metabolism, suggesting that reduced energetic resilience may link cardiometabolic stress to cardiac dysfunction.

These transcriptomic and lipidomic findings reinforce the rationale for therapies targeting metabolic adaptation, mitochondrial function and cellular energetics in HFpEF. Several genes encoding proteins involved in mitophagy and mitochondrial maintenance were suppressed, including *Pink1, Mfn2* and *Samm50*, together with induction of oxidative stress defence pathways.^47–49^ Reduced mitochondrial quality-control capacity combined with increased oxidative stress suggests impaired energy pathways and reduced mitochondrial function.^44^ These changes occurred alongside activation of inflammatory pathways and altered calcium-handling programs, reinforcing the concept that the combination of energetic dysfunction, oxidative stress and inflammation induce HFpEF.^16 2^ These transcriptional changes are consistent with impaired mitochondrial quality control and support a link between endothelial dysfunction, energetic stress and activation of injury-associated signalling pathways.

The coordinated suppression of genes involved in calcium handling and sarcoplasmic reticulum function represents another important finding. Reduced expression of *Ryr2, Hrc* and *Trdn* suggests disruption of intracellular calcium cycling, while suppression of Kcnip2, Kcnd2 and Trpm4 is consistent with electrical remodelling and altered excitation-contraction coupling.^21,23^ These abnormalities were present in both female treatment groups, suggesting that dysregulation of calcium homeostasis represents a fundamental myocardial response to cardiometabolic stress rather than a feature of advanced disease.^50^ Because calcium cycling directly influences ventricular relaxation and myocardial reserve, suppression of these pathways may contribute to the development of diastolic dysfunction observed in the current model. Similar mechanisms have been reported in both HFD/L-NAME models and clinical HFpEF studies.^14,51^ We also identified regulation of genes associated with circadian rhythm and the MR and GR.^52,53^ We and others have also shown disruption of the circadian clock in diabetes and metabolic disease^54,55^ suggesting that circadian and corticosteroid-responsive transcriptional networks may contribute to the cardiac response to chronic cardiometabolic stress in female hearts.

Lipidomic profiling of serum demonstrated both conserved and sex-specific responses to chronic cardiometabolic stress.^56,57^ While broad alterations in sphingolipid and phospholipid pathways were observed in both sexes, important differences emerged among specific phosphoinositol and lysophospholipid species. Despite marked differences in cardiac phenotype, male and female mice exhibited similar regulation of several circulating lipid classes, suggesting that systemic metabolic responses to HFD and L-NAME were largely conserved across groups.

Recent studies have highlighted important roles for altered lipid utilisation, ketone metabolism and mitochondrial substrate selection in HFpEF pathogenesis.^26,29,58,59^ Ceramides, HexCer and Hex2Cer were increased in multiple treatment groups, consistent with previous studies linking activation of sphingolipid pathways to obesity, insulin resistance and cardiovascular disease. Because similar sphingolipid changes were observed in groups that developed different cardiac phenotypes, these pathways are more likely to reflect exposure to cardiometabolic stress than the specific form of HF that subsequently developed.^29,56^

Our finding of increased circulating acylcarnitines in treated groups is consistent with altered fatty acid metabolism and mitochondrial substrate utilisation. Acylcarnitines accumulate when mitochondrial fatty acid oxidation is incomplete and have been associated with impaired metabolic flexibility and mitochondrial dysfunction in obesity, diabetes and HFpEF.^60,61^ Together with the transcriptomic evidence of mitochondrial pathway regulation, the acylcarnitine profile is consistent with altered myocardial energy metabolism in a number of tissues in response to the HFD. Similarly, regulation of plasmalogen species may reflect adaptive remodelling of membrane lipid composition and antioxidant responses.^62^ Plasmalogens have important roles in maintaining membrane integrity and protecting against oxidative stress, and increased levels in treated animals may represent a compensatory response to increased oxidative and metabolic stress.^63^

One of the clearest sex-specific lipid responses involved PI and LPC species, which regulated differently in male and female mice. Phosphoinositols are critical mediators of intracellular signalling pathways regulating insulin sensitivity, cellular energetics and membrane dynamics, whereas LPC species have recognised roles in inflammatory signalling, endothelial dysfunction and lipid turnover.^62,63^ These findings support a model in which systemic lipidomic remodelling reflects exposure to cardiometabolic stress, myocardial metabolic adaptation, which involves mitochondrial remodelling, may determine functional cardiac outcomes.

Myocardial fibrosis is frequently regarded as a hallmark of HFpEF; however, neither male nor female mice demonstrated substantial increases in collagen deposition despite cardiac functional changes and substantial regulation of the transcriptome. Recent studies seeking to optimise the HFpEF phenotype in mouse models reported cardiac fibrosis in prolonged HFD/L-NAME or DOCP/salt-based HFpEF models in both male and female mice.^18^ Although extracellular matrix pathways were activated at the transcriptional level in our study in female mice, measurable increases in tissue collagen was not seen at the 15-week study endpoint in both male and female hearts. These findings suggest that fibrosis is not an obligatory early feature of HFpEF-like remodelling and may be detected at later timepoints, with greater hypertensive burden, or additional profibrotic stimuli such as mineralocorticoid/salt activation.^18,64,65^ Metabolic dysfunction, calcium-handling abnormalities, loss of NO signalling and energetic stress may therefore precede overt structural remodelling and represent earlier phases of disease development. These data are consistent with clinical evidence of variability in tissue fibrosis among patients with HFpEF and highlights that non-fibrotic mechanisms contribute to disease pathogenesis in HFpEF.^16,66^

We propose that the present findings are potentially relevant to evaluating changes to HFpEF therapies. The clinical benefits observed with SGLT2 inhibitors, GLP-1 receptor agonists and finerenone suggest that activation of metabolic, inflammatory and cardiorenal pathways is occurs in many patients with HFpEF.^67–71^, Our data extend these observations by indicating that NOS inhibition and metabolic adaptation can influence the cardiac response to cardiometabolic stress. Rather than identifying a single disease mechanism, these therapies support a model in which multiple biological pathways contribute to tissue pathology. In line with clinical observations, our data suggest that preservation of nitric oxide signalling and metabolic flexibility determine whether cardiometabolic stress remains compensated, induces HFpEF-like remodelling or progresses towards myocardial injury and systolic dysfunction. Consequently, interventions targeting mitochondrial function, endothelial adaptation, calcium handling and inflammatory signalling may modify tissue remodelling before irreversible dysfunction develops.

It is important to note that we did not evaluate a lower dose of L-NAME in male mice and therefore cannot determine whether males also exhibit a HFpEF-like phenotype with a lower dose of L-NAME. The present study design focused on endpoint phenotypes and cannot establish the temporal sequence linking tissue remodelling pathways to systolic dysfunction. We propose that this point will be important to analyse in future studies. Phosphatidylcholine species were quantified but not included in the final lipidomic analyses because the dominant changes reflected differences in dietary lipid sources between chow and high-fat diets rather than cardiac phenotype. Excluding these species reduced diet-driven effects and allowed greater focus on lipid pathways associated with the response to cardiometabolic stress. Finally, longer-duration studies will be required to determine whether fibrosis and advanced structural remodelling emerge with continued cardiometabolic stress.

## Conclusions

Our data support existing evidence that NOS inhibition is important for driving HFpEF-like remodelling. Rather than promoting progressively more severe HFpEF, increasing NOS inhibition redirected the myocardium towards inflammatory, mitochondrial and injury-associated responses. This observation supports the concept that HFpEF develops within a specific range of NO signalling and raises the possibility that preserving endothelial health is particularly important during the earliest stages of disease development. Taken together, our findings suggest that future HFpEF models should move beyond reproducing a single endpoint phenotype and instead investigate mechanisms governing transitions between adaptive remodelling, HFpEF and systolic failure. Future studies should define whether a true endothelial stress threshold exists, determine the mechanisms underlying female protection and establish how mitochondrial dysfunction interacts with calcium handling and inflammatory signalling. Evaluating whether metabolic therapies alter myocardial pathogenic changes at different stages of the disease has the potential to provide insights for precision approaches to HFpEF that account for biological sex, NOS inhibition and metabolic phenotype.

## What’s New

- Biological sex and NOS inhibition determine different functional outcomes of cardiometabolic HF.
- HFpEF-like remodelling develops in female mice exposed to intermediate endothelial stress, whereas greater NOS inhibition promoted inflammatory and injury-associated responses.
- Similar cardiometabolic stress produced fundamentally different cardiac phenotypes despite broadly conserved systemic lipidomic remodelling.

HFpEF is often viewed as a progressive consequence of cardiometabolic disease. Our findings suggest that biological sex and NOS inhibition influence whether cardiometabolic stress results in adaptive remodelling, HFpEF-like dysfunction or systolic HF. These data provide a potential explanation for the marked heterogeneity observed among patients with similar cardiometabolic risk profiles and suggest that therapies targeting endothelial dysfunction may influence the cellular and tissue pathways to disease rather than simply disease severity.

## Acknowledgments

Melissa Lyttle for assistance with blood pressure measurements.

## Data Availability

RNA sequencing data, mass spectrometry lipidomics data and additional data supporting the findings of this study are available from the corresponding author upon reasonable request.

## References

1. Wu, J., et al. Artificial intelligence methods for improved detection of undiagnosed heart failure with preserved ejection fraction. Eur J Heart Fail 26, 302–310 (2024).

2. Peikert, A., Vacca, A., Norata, G.D., Schiattarella, G.G. & Osto, E. Cellular Interactions and Immunometabolic Mechanisms in Heart Failure With Preserved Ejection Fraction: From Molecular Mechanisms to Clinical Evidence. Circ Heart Fail 19, e012674 (2026).

3. Shibutani, Y., Aoki, F., Imaoka, T., Suzuki, A. & Tajiri, K. Clinical Characteristics and Outcomes of Heart Failure with Preserved Ejection Fraction Versus Reduced Ejection Fraction in Patients Receiving Anticancer Drugs. Cardiovasc Toxicol 26(2026).

4. van Essen, B.J., et al. Obesity and inactivity cluster the strongest risk factor for the development of heart failure in a population-based study. Int J Cardiol 442, 133914 (2026).

5. Alexandrou, M., Parikh, N.I., Brilakis, E.S., Lloyd-Jones, D.M. & Xanthakis, V. Insights Into Cardiovascular Disease in Women From the Framingham Heart Study. JACC Adv 5, 102722 (2026).

6. Kaur, G. & Lau, E. Sex differences in heart failure with preserved ejection fraction: From traditional risk factors to sex-specific risk factors. Womens Health (Lond) 18, 17455057221140209 (2022).

7. Kapelios, C.J., Shahim, B., Lund, L.H. & Savarese, G. Epidemiology, Clinical Characteristics and Cause-specific Outcomes in Heart Failure with Preserved Ejection Fraction. Card Fail Rev 9, e14 (2023).

8. Handford, C., et al. Targeting Cardiac Metabolism in Heart Failure with PPARalpha Agonists: A Review of Preclinical and Clinical Evidence. Biomedicines 13(2025).

9. Reimer Jensen, A.M., et al. Integrated trajectories of systolic and diastolic function differentially associate with risk for heart failure with preserved and reduced ejection fraction and proteomic profiles. Eur J Heart Fail 27, 2906–2917 (2025).

10. Lund, L.H., et al. Outcomes of heart failure with reduced, mildly reduced, or preserved ejection fraction: the ESC HF III registry. Eur Heart J 47, 2326–2341 (2026).

11. Gao, S., et al. Animal models of heart failure with preserved ejection fraction (HFpEF): from metabolic pathobiology to drug discovery. Acta Pharmacol Sin 45, 23–35 (2024).

12. Daou, D., Tong, D., Schiattarella, G.G., Gillette, T.G. & Hill, J.A. What Is Cardiometabolic HFpEF and How Can We Study it Preclinically? JACC Basic Transl Sci 10, 101295 (2025).

13. Ostrominski, J.W., et al. Performance of the HFpEF-ABA, H(2)FPEF, and HFA-PEFF Algorithms in Heart Failure With Preserved Ejection Fraction: A Participant-Level Pooled Analysis of Randomized Clinical Trials. J Card Fail (2026).

14. Schiattarella, G.G., et al. Nitrosative stress drives heart failure with preserved ejection fraction. Nature 568, 351–356 (2019).

15. Withaar, C., Lam, C.S.P., Schiattarella, G.G., de Boer, R.A. & Meems, L.M.G. Heart failure with preserved ejection fraction in humans and mice: embracing clinical complexity in mouse models. Eur Heart J 42, 4420–4430 (2021).

16. Adrah, Y., et al. Defining HFpEF in rodents: a systematic review. Cardiovasc Res 121, 2134–2143 (2025).

17. Roh, J., Hill, J.A., Singh, A., Valero-Munoz, M. & Sam, F. Heart Failure With Preserved Ejection Fraction: Heterogeneous Syndrome, Diverse Preclinical Models. Circ Res 130, 1906–1925 (2022).

18. McIntosh, B., et al. Optimized murine HFpEF models for translational preclinical studies. ESC Heart Fail 13(2026).

19. Abubakar, M., et al. Sex-specific differences in risk factors, comorbidities, diagnostic challenges, optimal management, and prognostic outcomes of heart failure with preserved ejection fraction: A comprehensive literature review. Heart Fail Rev 29, 235–256 (2024).

20. Brouwers, F.P., et al. Incidence and epidemiology of new onset heart failure with preserved vs. reduced ejection fraction in a community-based cohort: 11-year follow-up of PREVEND. Eur Heart J 34, 1424–1431 (2013).

21. Bienvenu, L.A., Morgan, J., Reichelt, M.E., Delbridge, L.M.D. & Young, M.J. Chronic in vivo nitric oxide deficiency impairs cardiac functional recovery after ischemia in female (but not male) mice. J Mol Cell Cardiol 112, 8–15 (2017).

22. Methawasin, M., et al. An ovary-intact postmenopausal HFpEF mouse model; menopause is more than just estrogen deficiency. Am J Physiol Heart Circ Physiol 328, H719–H733 (2025).

23. Bienvenu, L.A., et al. Cardiomyocyte Mineralocorticoid Receptor Activation Impairs Acute Cardiac Functional Recovery After Ischemic Insult. Hypertension 66, 970–977 (2015).

24. Kanki, M., et al. Balcinrenone Shows a Unique Regulation of Potassium Excretion in Streptozotocin-induced Diabetes in Male Mice. Endocrinology 167(2026).

25. Poole, D.C., et al. Guidelines for animal exercise and training protocols for cardiovascular studies. Am J Physiol Heart Circ Physiol 318, H1100–H1138 (2020).

26. Bond, S.T., et al. Mitochondrial damage in muscle specific PolG mutant mice activates the integrated stress response and disrupts the mitochondrial folate cycle. Nat Commun 16, 2338 (2025).

27. Bond, S.T., et al. Tissue-specific expression of Cas9 has no impact on whole-body metabolism in four transgenic mouse lines. Mol Metab 53, 101292 (2021).

28. Wearing, O.H., et al. Guidelines for assessing ventricular pressure-volume relationships in rodents. Am J Physiol Heart Circ Physiol 328, H120–H140 (2025).

29. Belkin, T.G., et al. An optimized plasmalogen modulating dietary supplement provides greater protection in a male than female mouse model of dilated cardiomyopathy. J Mol Cell Cardiol Plus 11, 100273 (2025).

30. Huynh, K., et al. High-Throughput Plasma Lipidomics: Detailed Mapping of the Associations with Cardiometabolic Risk Factors. Cell Chem Biol 26, 71–84 e74 (2019).

31. Heanue, S., et al. Temporal mineralocorticoid receptor activation regulates the molecular clock and transcription of cardiovascular disease modulators in myeloid cells. Am J Physiol Heart Circ Physiol 328, H1318–H1332 (2025).

32. Rickard, A.J., et al. Deletion of mineralocorticoid receptors from macrophages protects against deoxycorticosterone/salt-induced cardiac fibrosis and increased blood pressure. Hypertension 54, 537–543 (2009).

33. Armanini, D., Sabbadin, C., Dona, G., Clari, G. & Bordin, L. Aldosterone receptor blockers spironolactone and canrenone: two multivalent drugs. Expert Opin Pharmacother 15, 909–912 (2014).

34. Team, R.C. R: A language and environment for statistical computing. R Foundation for Statistical Computing.. (https://www.R-project.org/, 2024).

35. Shah, S.J., et al. Phenomapping for novel classification of heart failure with preserved ejection fraction. Circulation 131, 269–279 (2015).

36. Obokata, M., Reddy, Y.N.V., Pislaru, S.V., Melenovsky, V. & Borlaug, B.A. Evidence Supporting the Existence of a Distinct Obese Phenotype of Heart Failure With Preserved Ejection Fraction. Circulation 136, 6–19 (2017).

37. Obokata, M., et al. Ventricular-Arterial Coupling and Exercise-Induced Pulmonary Hypertension During Low-Level Exercise in Heart Failure With Preserved or Reduced Ejection Fraction. J Card Fail 23, 216–220 (2017).

38. Actis Dato, V., Lange, S. & Cho, Y. Metabolic Flexibility of the Heart: The Role of Fatty Acid Metabolism in Health, Heart Failure, and Cardiometabolic Diseases. Int J Mol Sci 25(2024).

39. Aryal, A., et al. Cardiac metabolic remodeling drives dicarbonyl stress-induced mitochondrial dysfunction in experimental heart failure with preserved ejection fraction. Am J Physiol Heart Circ Physiol 330, H1689–H1701 (2026).

40. Ganthan, R.R. & Rotatori, F. Women in Heart Failure With Preserved Ejection Fraction Clinical Trials: Bridging the Gap Between Trial Enrollment and Real-World Disease Burden. Cardiol Rev (2026).

41. Ritterhoff, J., et al. Increasing fatty acid oxidation elicits a sex-dependent response in failing mouse hearts. J Mol Cell Cardiol 158, 1–10 (2021).

42. Gambardella, J., Lombardi, A. & Santulli, G. Metabolic Flexibility of Mitochondria Plays a Key Role in Balancing Glucose and Fatty Acid Metabolism in the Diabetic Heart. Diabetes 69, 2054–2057 (2020).

43. Shehadeh, L.A., Robleto, E. & Lopaschuk, G.D. Cardiac energy substrate utilization in heart failure with preserved ejection fraction: reconciling conflicting evidence on fatty acid and glucose metabolism. Am J Physiol Heart Circ Physiol 328, H1267–H1295 (2025).

44. Koay, Y.C., et al. The Heart Has Intrinsic Ketogenic Capacity that Mediates NAD(+) Therapy in HFpEF. Circ Res 136, 1113–1130 (2025).

45. O’Sullivan, J.F., et al. Cardiac Substrate Utilization and Relationship to Invasive Exercise Hemodynamic Parameters in HFpEF. JACC Basic Transl Sci 9, 281–299 (2024).

46. Taegtmeyer, H., Wilson, C.R., Razeghi, P. & Sharma, S. Metabolic energetics and genetics in the heart. Ann N Y Acad Sci 1047, 208–218 (2005).

47. Zhou, Y., et al. Mitochondrial outer membrane protein Samm50 protects against hypoxia-induced cardiac injury by interacting with Shmt2. Cell Signal 120, 111219 (2024).

48. Zhang, H., et al. PINK1 modulates Prdx2 to reduce lipotoxicity-induced apoptosis and attenuate cardiac dysfunction in heart failure mice with a preserved ejection fraction. Clin Transl Med 15, e70166 (2025).

49. Zhang, S., et al. SGLT2 inhibitor dapagliflozin treats heart failure with preserved ejection fraction via the SIRT1/PGC-1alpha pathway. Acta Biochim Biophys Sin (Shanghai) xx, 1624–1636 (2026).

50. Morciano, G., et al. Calcium dysregulation in heart diseases: Targeting calcium channels to achieve a correct calcium homeostasis. Pharmacol Res 177, 106119 (2022).

51. Elbassioni, A.A.M., et al. Targeting RUNX1 protects against diastolic dysfunction in a two-hit mouse model of heart failure with preserved ejection fraction. Cardiovasc Res 122, 1318–1328 (2026).

52. Rickard, A.J., et al. Cardiomyocyte mineralocorticoid receptors are essential for deoxycorticosterone/salt-mediated inflammation and cardiac fibrosis. Hypertension 60, 1443–1450 (2012).

53. Fletcher, E.K., et al. Deoxycorticosterone/Salt-Mediated Cardiac Inflammation and Fibrosis Are Dependent on Functional CLOCK Signaling in Male Mice. Endocrinology 158, 2906–2917 (2017).

54. Young, M.E., Wilson, C.R., Razeghi, P., Guthrie, P.H. & Taegtmeyer, H. Alterations of the circadian clock in the heart by streptozotocin-induced diabetes. J Mol Cell Cardiol 34, 223–231 (2002).

55. Kanki, M. & Young, M.J. The mineralocorticoid receptor: a new chapter for therapeutic regulation of diabetic cardiomyopathy. J Mol Cell Cardiol 213, 1–13 (2026).

56. Tham, Y.K., et al. Lipidomic Profiles of the Heart and Circulation in Response to Exercise versus Cardiac Pathology: A Resource of Potential Biomarkers and Drug Targets. Cell Rep 24, 2757–2772 (2018).

57. Tham, Y.K., et al. Distinct lipidomic profiles in models of physiological and pathological cardiac remodeling, and potential therapeutic strategies. Biochim Biophys Acta Mol Cell Biol Lipids 1863, 219–234 (2018).

58. Koay, Y.C., et al. Plasma levels of trimethylamine-N-oxide can be increased with ‘healthy’ and ‘unhealthy’ diets and do not correlate with the extent of atherosclerosis but with plaque instability. Cardiovasc Res 117, 435–449 (2021).

59. Wang, Y.C., et al. Indole-3-Propionic Acid Protects Against Heart Failure With Preserved Ejection Fraction. Circ Res 134, 371–389 (2024).

60. Mihalik, S.J., et al. Increased levels of plasma acylcarnitines in obesity and type 2 diabetes and identification of a marker of glucolipotoxicity. Obesity (Silver Spring) 18, 1695–1700 (2010).

61. Bernardo, B.C., et al. Lipidomic Profiling of a Preclinical Model of Streptozotocin-Induced Diabetic Cardiomyopathy Reveals Potential Plasma Biomarkers. Heart Lung Circ 34, 739–742 (2025).

62. Schooneveldt, Y.L., et al. Alterations in ether lipid metabolism in obesity revealed by systems genomics of multi-omics datasets. PLoS Biol 23, e3003349 (2025).

63. Pradas, I., et al. Lipidomics Reveals a Tissue-Specific Fingerprint. Front Physiol 9, 1165 (2018).

64. Li, S., et al. Nicotinamide-N-methyltransferase inhibition improves cardiac function and structure in a heart failure with preserved ejection fraction mouse model. Pharmacol Res 217, 107820 (2025).

65. Meral, D., et al. Reversible Fibroblast Trajectories Regulated by MR Underlie Diastolic Dysfunction. Circ Res 138, e327301 (2026).

66. Paulus, W.J. & Tschope, C. A novel paradigm for heart failure with preserved ejection fraction: comorbidities drive myocardial dysfunction and remodeling through coronary microvascular endothelial inflammation. J Am Coll Cardiol 62, 263–271 (2013).

67. Anker, S.D., et al. Efficacy of empagliflozin in heart failure with preserved versus mid-range ejection fraction: a pre-specified analysis of EMPEROR-Preserved. Nat Med 28, 2512–2520 (2022).

68. Solomon, S.D., et al. Dapagliflozin in Heart Failure with Mildly Reduced or Preserved Ejection Fraction. N Engl J Med 387, 1089–1098 (2022).

69. Solomon, S.D., et al. Finerenone in Heart Failure with Mildly Reduced or Preserved Ejection Fraction. N Engl J Med 391, 1475–1485 (2024).

70. Ostrominski, J.W., et al. Efficacy and Safety of Finerenone in Heart Failure With Preserved Ejection Fraction: A FINE-HEART Analysis. JACC Heart Fail 13, 102497 (2025).

71. Packer, M., et al. Tirzepatide for Heart Failure with Preserved Ejection Fraction and Obesity. N Engl J Med 392, 427–437 (2025).

